# BSAlign: a library for nucleotide sequence alignment

**DOI:** 10.1101/2024.01.15.575791

**Authors:** Haojing Shao, Jue Ruan

**Author notes:** Contributed equally.

## Abstract

Increasing the accuracy of the nucleotide sequence alignment is an essential issue in genomics research. Although classic dynamic-programming algorithms (e.g., Smith-Waterman and Needleman–Wunsch) guarantee to produce the optimal result, their time complexity hinders the application of large-scale sequence alignment. Many optimization efforts that aim to accelerate the alignment process generally come from three perspectives: re-designing data structures (e.g., diagonal or striped Single Instruction Multiple Data (SIMD) implementations), increasing the number of parallelisms in SIMD operations (e.g., difference recurrence relation), or reducing searching space (e.g., banded dynamic programming). However, no methods combine all these three aspects to build an ultra-fast algorithm. We have developed a Banded Striped Aligner(library) named BSAlign that delivers accurate alignment results at an ultra-fast speed by knitting a series of novel methods together to take advantage of all of the aforementioned three perspectives with highlights such as active F-loop in striped vectorization and striped move in banded dynamic programming. We applied our new acceleration design on both regular and edit-distance pairwise alignment. BSAlign achieved 2-fold speed-up than other SIMD based implementations for regular pairwise alignment, and 1.5 to 4-fold speedup in edit distance based implementations for long reads. BSAlign is implemented in C programing language and is available at https://github.com/ruanjue/bsalign.

## 1. Introduction

Nucleotide sequence alignment is a way to arrange and compare DNA/RNA sequences from different sources to identify their regions of similarity. Two classic algorithms, namely Needleman-Wunsch algorithm[1] and Smith-Waterman algorithm[2], are commonly used for sequence alignment. They handle sequence alignment by solving a dynamic programming problem in which a scoring matrix is calculated and an optimal path from the cell with a maximal score is returned. Although these two methods have shown high capability in finding optimal alignment results, they require quadratic time complexity and rapidly degenerate especially when processing long sequences. To accelerate the alignment process, three major categories of optimization techniques have been developed along the way.

### Single Instruction Multiple Data (SIMD)

The first optimization category is to re-design the data structure of the scoring matrix calculation to resolve data dependencies between neighboring cells so that the conditional branch within the inner loop of the dynamic programming algorithm can be eliminated and hence more efficient in parallelization techniques such as SIMD. Among the initial trials in this category, Wozniak[3] has presented an implementation to store values parallel to the minor diagonal to eliminate the conditional branch in the inner loop of traditional implementation and achieved a 2x speedup. In a different trial, Rognes et al.[4] introduced another implementation to store values parallel to the query sequences. Compared to Wozniak’s implementation, an advantage of Rognes’s design is that it only needs to compute the query profile once for the entire reference sequences. However, the disadvantage is that conditional branches are placed in the inner loop when evaluating F matrix. The length of a single instruction ranges from 128-bit to 512-bit for recent tools such as BGSA[5], SeqAn[6] and AnySeq[7].

### Striped SIMD and F evaluation

To combine the merits of both Wozniak[3] and Rognes[4], Farrar[8] fixed these disadvantages by introducing a layout of query sequences that are parallel to the SIMD registers but are accessed in a striped pattern, which only computes query profile once and moves the conditional F matrix evaluation outside of the inner loop. As a result, Farrar’s striped vectorization successfully speeds up the Smith-Waterman algorithm and has been adopted by many aligners, such as BWA-SW[9], Bowtie2[10], and SSW library[11]. However, cells in the same register are not always independent of each other. Farrar[8] solved this problem by adding a correction loop for every F element, which may iterate many times when the indels are long enough.

### Difference recurrence relation

The next optimization category is to increase the number of parallelisms in SIMD operations such as difference recurrence relation [12]. Since the traditional pairwise alignment stores and calculates the absolute values in the score matrix, it limits the width of SIMD operation as the sequence length increase. The difference recurrence relation solves this problem by only storing and calculating the differences between the adjacent cells, which keeps the full width of SIMD operation regardless of the sequence length. For example, the number of bits for storing the absolute value of a single cell is 16 or even 32. But it can reduce to 8 bits if just storing the differences between cells. Therefore, the number of parallelisms increases by 2 to 4 times.

### Banded dynamic programming

Another optimization category is reducing the search space such as banded dynamic programming(DP). Instead of calculating the whole score matrix, banded DP maintains a hypothetical “band” around cells with maximal scores and only calculates the scores for cells within the “band” and skips calculating the remaining cells within the matrix[13, 14]. How to combine the method of using SIMD (minor diagonal or striped) and the idea of reducing the search space is not clear, especially when the input sequences contain abundant indel errors by third-generation sequencers. Suzuki et al.[14] proposed a minor diagonal SIMD adaptive banded DP algorithm, which is implemented and improved in a popular long read mapper minimap2[15]. Since the striped SIMD method[8] proved to be six times faster than the minor diagonal and other SIMD method[3] in SW algorithm without banded DP, the algorithm to combine the best SIMD method with banded DP is not developed yet.

### Block aligner and wavefront algorithm

Recent methods block aligner(BA)[16] and wavefront algorithm(WFA)[17] manage to reduce the search space around the diagonal by two innovative approaches. Block aligner starts the alignment by a small square block and extends the block dynamically until the endpoint is reached. The block could be shifted either down or right according to the sum of the cells. The size of the extended block may double when a Y-drop condition is met. The width of the block (band) depends on the sequence’s identity. Unlike the block aligner, the wavefront algorithm regards the global alignment as the wave spreading from the start point to the endpoint. WFA extends the wave step-by-step until the end-point is reached. To speed up, WFA utilizes homologous region between the sequences to skip the path(wavefront) that are unlikely to lead to the optimal solution. The search space for the wavefront aligner is wave-like banding along the diagonal.

Overall, we present a new library with an aligner, BSAlign, that is able to combine merits from the aforementioned optimization techniques without bringing their respective limitations. Firstly, we developed an active F loop evaluation algorithm in the striped vectorization[8] to reduce the redundant F matrix recalculation, which accelerates the evaluation of the scoring matrix. We also introduced difference recurrence relation and developed a banded DP striped move algorithm to efficiently combine the striped SIMD method and banded DP. Finally, we designed a fast bit-vector algorithm to further speed-up edit distance based alignment.

## 2. Materials and Methods

### 2.1. Overview

We developed a set of new methods to address the pairwise sequence alignment problem by adopting advantages from previous work like striped vectorization [8], difference recurrence relation[12], banded dynamic programming[13], and by proposing novel improvements like a technique called active F-loop evaluation, a set of newly derived recurrence relations, a variant of bit conversion for edit-distance alignment, and different levels of adjustments to integrate all the features into the BSAlign.

### 2.2. The global alignment of nucleotide sequences

In the beginning, the algorithm calculates the global alignment by the Needleman Wunsch algorithm[1]. The two sequences to be aligned, the query sequence and the reference sequence, are defined as Q and R. The length of the query sequence and reference sequence are then defined as *Q*_*len*_ and *R*_*len*_, respectively. A matching matrix S(*q*_*i*_, *r*_*j*_) is defined for all residue pairs (a, b) where *a, b* ∈ {*A, T, C, G}*. The matching score *S*(*q*_*i*_, *r*_*j*_) *<* 0 when *q*_*i*_! = *r*_*j*_ and *S*(*q*_*i*_, *r*_*j*_) *>* 0 when *q*_*i*_ == *r*_*j*_. The penalty for starting a gap and continuing a gap are defined as GapO (gap open, *GapO <* 0), GapE (gap extension, *GapE <* 0), and GapOE = GapO + GapE. We keep track of three scoring matrices: E, F, and H, where E represents the alignment score ending with a vertical gap, F represents the alignment score ending with a horizontal gap:

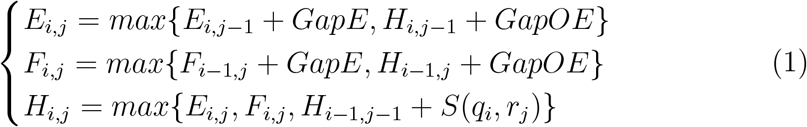

The cells for *H*_*i,j*_, *E*_*i,j*_ and *F*_*i,j*_ are filled by 0 when *i <* 1 or *j <* 1. In our implementation, we store *S*(*q*_*i*_, *r*_*j*_) in four query profile arrays: S(*Q*,A), S(*Q*,C), S(*Q*,T) and S(*Q*,G). We calculate the score matrix row by row and extract the *S*(*q*_*i*_, *r*_*j*_) from query profile column *S*(*Q, r*_*j*_). We simplify *S*(*q*_*i*_, *r*_*j*_) as *S*_*i,j*_ in this manuscript.

### 2.3. The striped SIMD data structure

To accelerate the pairwise alignment in the data structure, we first implemented striped SIMD[8] to the row of the score matrix as well as the query profile arrays. Assuming the query and reference sequences are the row and column in the score matrix, respectively. The row is divided into equal length segments, *S*. The number of segments, *p*, is equal to the number of cells being processed in a SIMD register. Take an example in 128 SSE. When processing byte integers (8-bit values) *p* = 16 and when processing word integers (16-bit values) *p* = 8. Hence, *p* is fixed in the algorithm and *S* depends on query length (or band width) *Q*_*len*_: *S* = ⌈*Q*_*len*_*/p*⌉(Fig1a). We first introduce the way to store the non-striped score matrix for each register *N* in the memory:

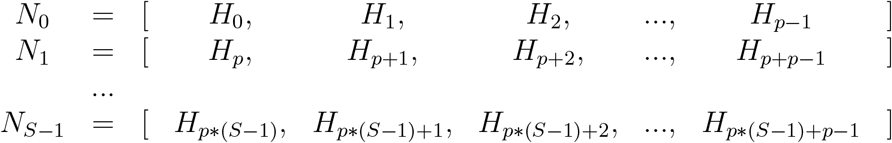

**Figure 1:**
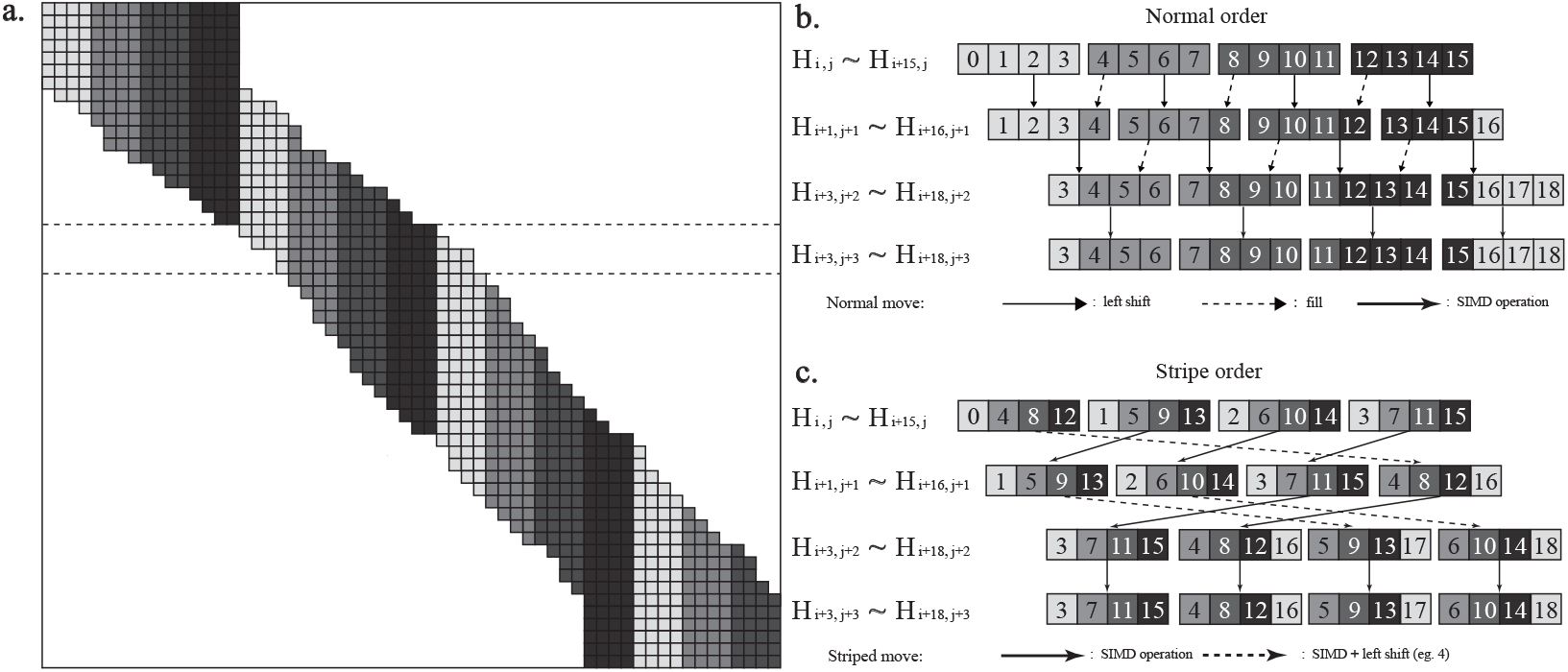
The striped move algorithm for each row. (**a**) Global visualization for the banded along the diagonal. (**b**) Detail example for row iteration in normal order. (**c**) Detail example for row iteration in striped order(striped move). Assuming the band width, the number of divided segments(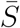) and the number of cells(*p*) in a register are 16, 4 and 4, respectively. In normal order, the cells are in the same color for the same register. Only the offset is numbered inside the cell.

The potential overflow cells in *N*_*S−*1_ are filled by minimum value. In the standard coordinate, there is an inner loop to compute *H* and *F* for each register.

After striped conversion, the memory will store the score matrix for each register *M* :

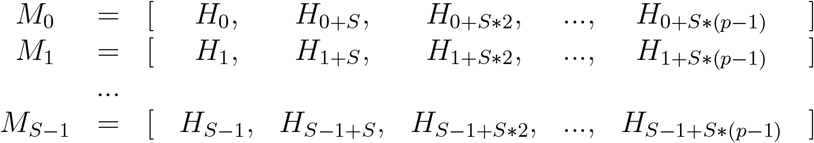

Hence, the equation to convert each value (*N*_*i,j*_) in non-striped SIMD(*N*) to any value (*M*_*i,j*_) in striped SIMD(*M*) is:

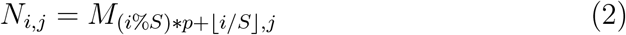

In the striped coordinate, all the inner loops to compute *H* and *F* are moved outside of the register. Now, *M*_*i*+1_ depends on *M*_*i*_. The initial one *M*_0_ is solved by the active F loop in the below subsection.

### 2.4. The striped move algorithm for banded DP

Another way to optimize the pairwise alignment is only focusing on the alignment along a diagonal band. A difficulty in applying banded DP to the striped SIMD method is that the entire striped SIMD data structure rearranges each time the band moves along the diagonal(Fig1b). To overcome this difficulty, we develop a method to move the striped SIMD data structure for banded DP(Fig1c).

In normal coordinate, the whole register stores(Fig1b):

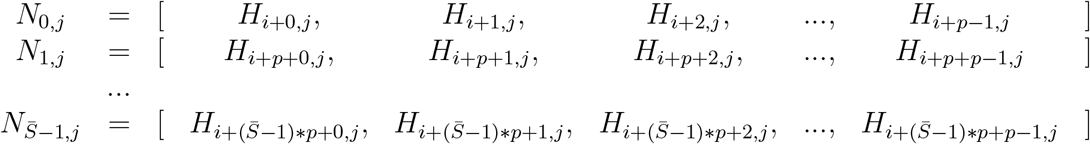

*i* and *j* are the first column and row coordinate in the memory and 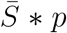 is the band width. In the banded DP, the movement of the current row indicates the selection of the optimal path. Moving zero, one and two cells to the right indicates one vertical gap, no gap and one horizontal gap, respectively(Fig1a). In our banded dynamic programming algorithm, we compare the *H* in the first and last register, move the current row zero, one or two cells to the right according to the comparison and prepare for the next row. In the next row, the start position *H*_*i*+*x,j*+1_ depends on the current row 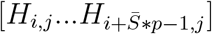 as following:

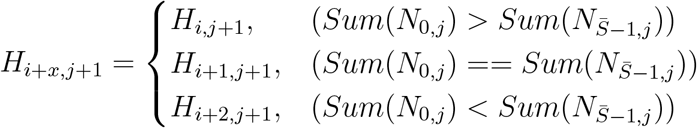

*H*_*i*+*x,j*+1_ move zero, one and two cell(s) to the right are showed as row j+3, j+1 and j+2 in Fig1b and Fig1c, respectively. We develop our striped move following the above equations. In striped coordinates, the whole register stores:

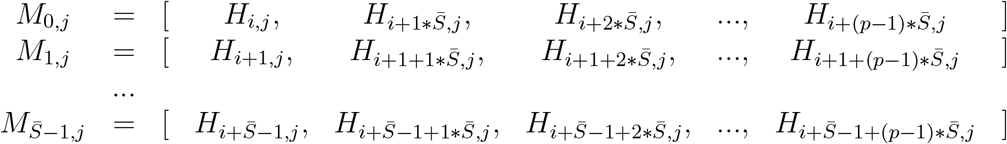

The memory stores all the register as 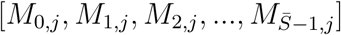. In the striped order, the cell order for the first register in j+1 row (such as [*H*_1,*j*+1_,*H*_5,*j*+1_,*H*_9,*j*+1_,*H*_13,*j*+1_]) is the same as the first, second and third register in the j row(such as [*H*_1,*j*_,*H*_5,*j*_,*H*_9,*j*_,*H*_13,*j*_]) for moving zero, one or two cells to the right, respectively(Fig1c). This holds true for all the registers except the last one or two registers. For the calculation of the next row, the memory is the following:

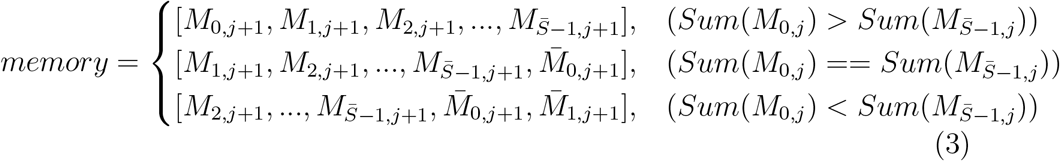

For the exception, the new 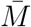 can be converted from the previous *M* (dash arrow in Fig1c, such as [*H*_1,*j*+1_,*H*_5,*j*+1_,*H*_9,*j*+1_,*H*_13,*j*+1_] to [*H*_5,*j*+2_,*H*_9,*j*+2_,*H*_13,*j*+2_,*H*_17,*j*+2_]). Take 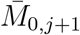 as an example:

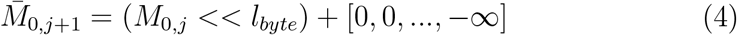

*l*_*byte*_ is the byte length of *H*. As the row moves one cell to the right, the whole cells move from 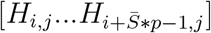 to 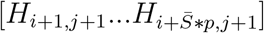. The addition cell 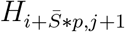 is set as the negative infinity and filled in the register. The negative infinity indicates this boundary cell will not be selected by equation1. The banded algorithm skips the calculation of boundary cells to speed up the global alignment. Using our striped move method, the whole striped SIMD data only needs bit operations to prepare for a new row in banded DP.

### 2.5. Difference Recurrence Relation

The third way to optimize the pairwise alignment is to increase the number of parallelisms (i.e., vector width) in SIMD operations. We choose to calculate the score matrix based on the difference recurrence relation[12] instead of the conventional stripped SIMD implementation[8]. We denote *h, e, f* as the relative score of the *H, E, F* score matrices respectively. We also define *u* matrix and *v* matrix to represent the vertical difference and horizontal difference within the *H* matrix, respectively.

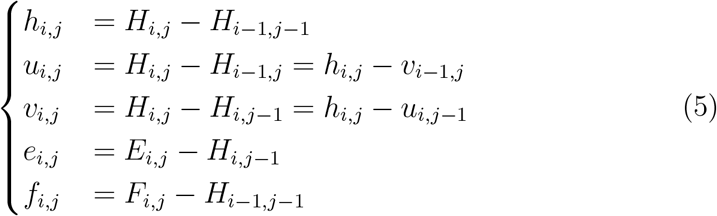

The definition of *e* and *f* matrix is asymmetry. Under the above definition, our difference recurrence relation can be expressed as:

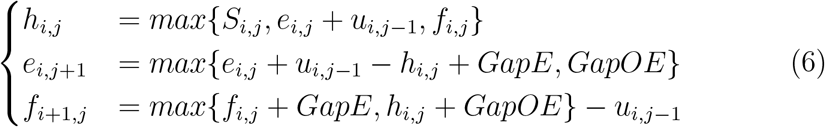

For every iteration in our implementation, we only store *u*_*i,j−*1_ and *e*_*i,j*_, and calculate *h*_*i,j*_, *v*_*i,j*_, *e*_*i,j*+1_ and *f*_*i*+1,*j*_ by equation 5 and 6.

### 2.6. Active F loop

For non-striped pairwise alignment, the horizontal score *F* and its difference *f* can be calculated step-by-step via previous cells. In striped order, some cells show up before their previous cells(*f*_4_, *f*_8_, *f*_12_ in Fig2). Thus, it raises a problem only in calculating the horizontal score *F* and its difference *f*. Farrar[8] initially developed a lazy F evaluation method to solve this problem(Fig2a). It corrected the value *F* via a couple of loops, which is time-consuming for sequences that contain long indel errors(Fig2d). In contrast to lazy F evaluation, we actively correct all the cells in advance and guarantee that the value of *H* is always corrected, providing a linear complexity solution to this problem in any situation(Fig2b,2e).

**Figure 2:**
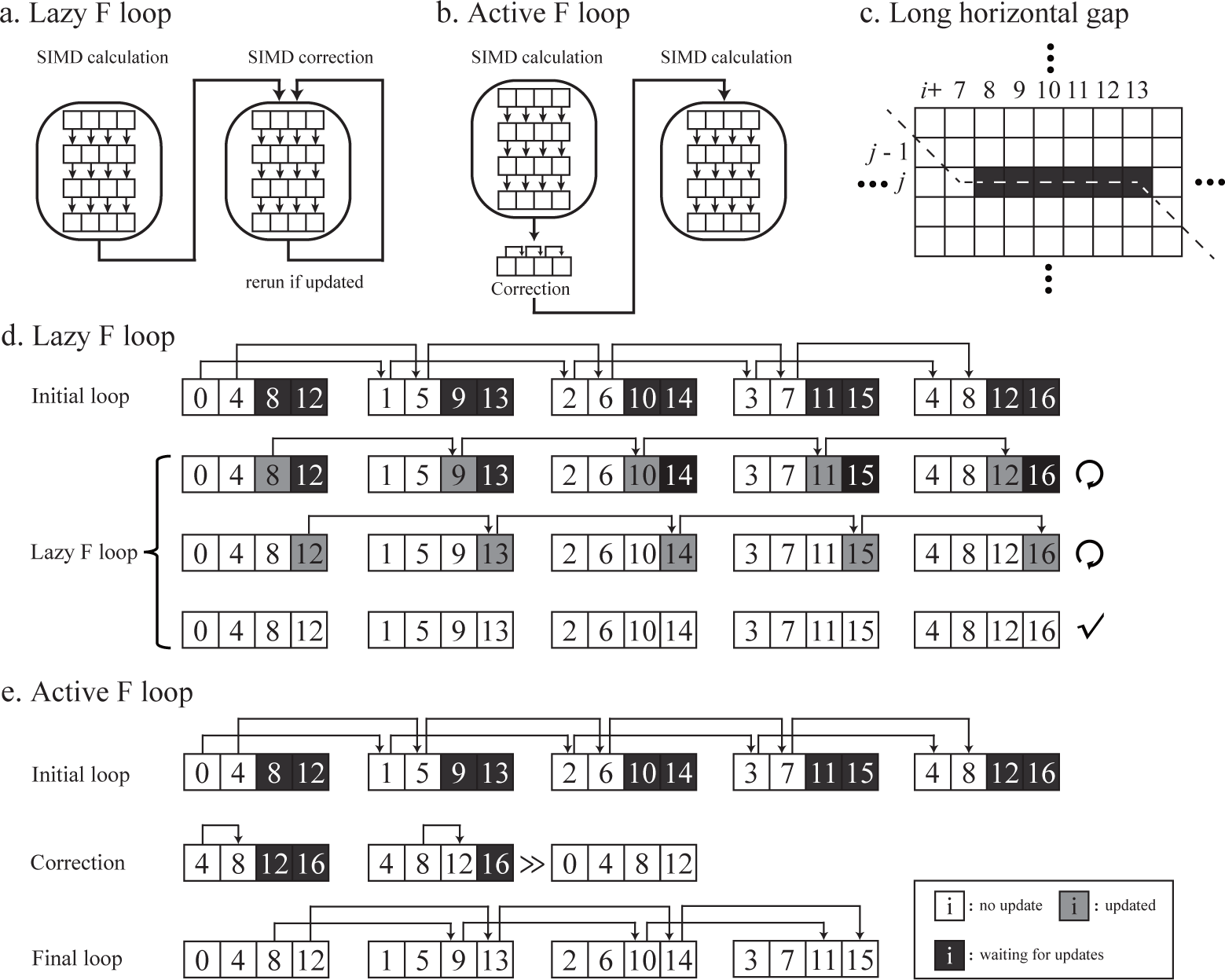
The lazy and active F loop algorithm inside a row. Assuming the band width, the number of divided segments(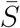) and the number of cells(*p*) in a register are 16, 4 and 4, respectively. (a) and (b) are the work flow for the lazy and active F loop, respectively. (c) is an example of long horizontal gap(*f*_7,*j*_ to *f*_13,*j*_). The dash line indicates the optimal path. (d) and (e) show how the lazy and active F loop solve the above example for each cell. Black box(waiting for updates) indicates the value is influenced by *F* and needs to be corrected. Grey box(updated) indicates the value is updated recently. White box(no update) indicates the value is the same as expected and correct. The arrows above digits indicate the first time being updated as the correct value. (d)*Initial loop*: All the cells in the first register are negative infinity. The algorithm calculates all the cells by standard SIMD calculation. *Lazy F loop*: The algorithm keeps correcting all the registers one by one until none of them is updated. This figure shows a situation that it takes 3 loops to guarantee that all the cells are corrected. (e)Unlike the lazy F loop, there is no difference between “updated” and “no update”. All the value is updated one times only. *Intital loop*: The same as the lazy F loop. *Correction*: Each cell in the first segment is checked by equation 7 and updated by the correct value. This extended register is also the first register (after the right shift) in the striped format. *Final loop*: When the first register is totally correct, the remaining segments are correct by SIMD calculation.

For most registers in the memory, *f* is smaller than *e* and *S*. The value of *H* does not source from *f* (FigS1a). Only the horizontal gap will *f* start to influence the value of *H*(FigS1b,c). In the initial loop, we set the negative infinity as the first register *MF*_0,*j*_ for horizontal difference *f* ([*f*_0_*f*_4_*f*_8_*f*_12_] in Fig2e):

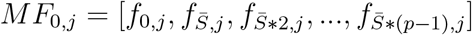

Because *f*_0,*j*_ will never contribute to *f*_1,*j*_, *f*_0,*j*_ (negative infinite) is always error-free(*f*_0_ in Fig2e top left). Then we calculate the whole matrix by equation 5 and 6. Because *f*_0,*j*_ is error-free, *f*_1,*j*_ to 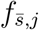 is correct(*f*_1_ to *f*_4_ in Fig2e). We save the last register 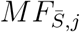:

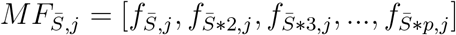

Note that we calculate *f*_*i*+1,*j*_ in equation 6, so 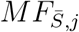 ([*f*_4_*f*_8_*f*_12_*f*_16_] in Fig2e) is the last register to store *f* instead of 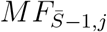. Because 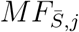 is calculated by 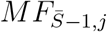, it solves the problems that 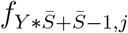 may update 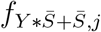 (*Y* = 0, 1, 2…, FigS1b).

The only exception is that the horizontal gap is long enough to penetrate all the 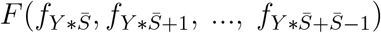 in the same cell position for all registers(Fig2c,S1c). If the F penetration happens, the value of *f*_*x,j*_ is smaller than 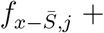 *gappenalty* (FigS1c). So we update the *f*_*x,j*_ as the corrected value in advance when we know F penetration has happened(correction in Fig2e). The equation to update all the *f* in the first register *MF*_0,*j*_ is following:

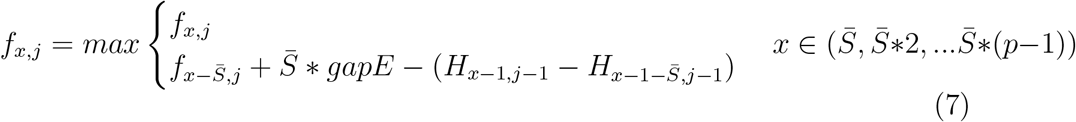

Now, the updated *f* solves the problems that 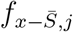 may update *f*_*x,j*_. So 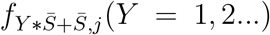 is corrected(*f*_8_,*f*_12_ in Fig2). We right shift the last register 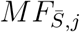 x bytes (length of *f*_0,*j*_) and update as the first register *MF*_0,*j*_([*f*_4_*f*_8_*f*_12_*f*_16_] to [*f*_0_*f*_4_*f*_8_*f*_12_] in Fig2). After the active F loop, we use the updated *f* as the initial value and recalculate all the values by equation 5,6(final loop in Fig2e). So the remaining values are corrected (*f*_5_,*f*_9_,*f*_13_,*f*_2_,*f*_6_,*f*_10_,*f*_7_,*f*_11_,*f*_15_ in Fig2 bottom). When all the values in *f* are corrected or error-free, all the values of *H* are corrected.

Specifically, the active F loop and the parallel scan in parasail[18, 19] are similar in general. One of the improvements is that the active F loop is implemented in difference recurrence relation, while the parallel scan[18] is implemented in the tradition way, storing and calculating the absolute values. Therefore, the active F loop can increase the number of parallelisms.

### 2.7. Edit distance

Calculating two sequences’ edit distance can be regarded as a special case of pairwise alignment when the mismatch and gap extend are both equal to 1 and, the match and gap open are both equal to 0. Since the difference between adjacent cells belongs to (−1,0,1), the number of bits for storing them is only 2. We can further increase the number of parallelisms using striped SIMD difference recurrence relation. As the number of bits decreases to 2, all the conditions can be enumerated. We converted the equation 5 and 6 to boolean logic to further accelerate the calculation.

To simplify the standard pairwise alignment, we only require *H, h, u* and *v*.

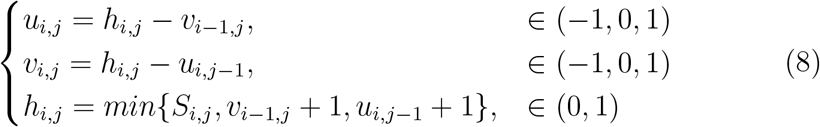

To minimize the computation resource, we defined a 2-bit binary code for boolean logic. For *h*_*i,j*_, *u*_*i,j*_ and *v*_*i,j*_, “-1,0,1” is converted to “10,00,01”. For *S*_*i,j*_, “0,1” is converted to “01,00”. All the conditions for calculating *h*_*i,j*_ from *S*_*i,j*_, *u*_*i,j−*1_ and *v*_*i−*1,*j*_ is enumerated as below(Table1). The new codes are inside the parentheses.

**Table 1:**
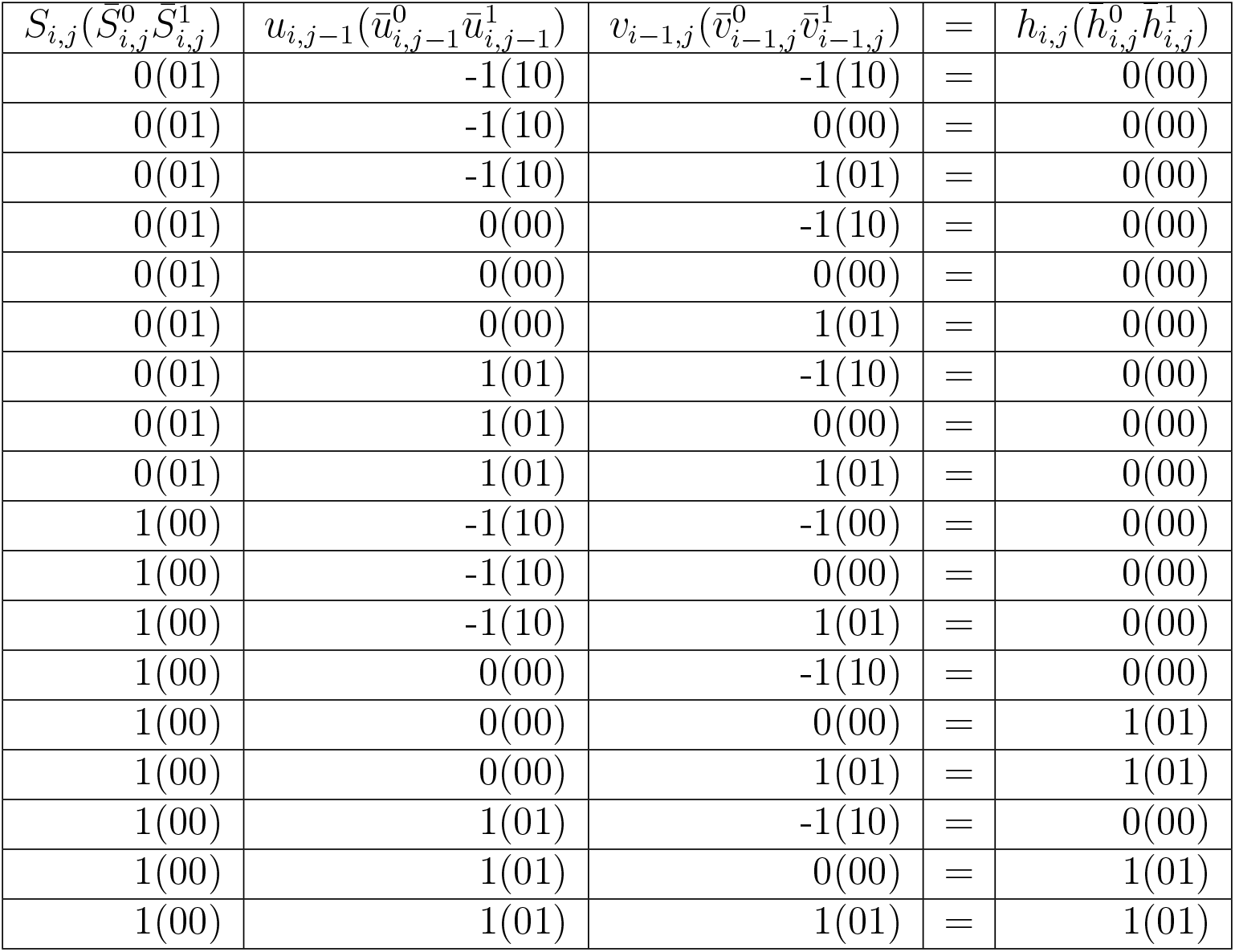
In edit distance mode, enumeration of conditions for converting *h*_*i,j*_ from *S*_*i,j*_, *u*_*i,j−*1_ and *v*_*i−*1,*j*_. The new binary codes are inside the parentheses.

Hence, the boolean logic for the new 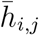 is following:

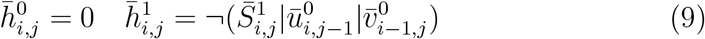

All the conditions for calculating *u*_*i,j*_ from *h*_*i,j*_ and *v*_*i−*1,*j*_ is enumerated as below(Table2).

**Table 2:** In edit distance mode, enumeration of conditions for converting *u*_*i,j*_ from *h*_*i,j*_ and *v*_*i−*1,*j*_. The new binary codes are inside the parentheses.

| $h_{i,j}(\bar{h}_{i,j}^0 \bar{h}_{i,j}^1)$ | $v_{i-1,j}(\bar{v}_{i-1,j}^0 \bar{v}_{i-1,j}^1)$ | = | $u_{i,j}(\bar{u}_{i,j}^0 \bar{u}_{i,j}^1)$ |
| --- | --- | --- | --- |
| 0(00) | 0(00) | = | 0(00) |
| 0(00) | 1(01) | = | -1(10) |
| 0(00) | -1(10) | = | 1(01) |
| 1(01) | 0(00) | = | 1(01) |
| 1(01) | 1(01) | = | 0(00) |

As *h*_*i,j*_ − *v*_*i−*1,*j*_ ∈ (0, 1), the condition of “*h*_*i,j*_ = 1 and *v*_*i−*1,*j*_ = −1” does not exist. Since *v*_*i,j*_ is symmetry to *u*_*i,j*_ in this definition, the boolean logic for 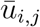 and 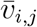 is following:

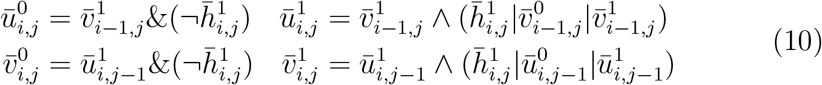

## 3. Experimental design

We implemented BSAlign with two modes: “align mode” for pairwise alignment by score matrix, and “edit mode” for pairwise alignment by minimum edit distance. For “align mode”, we compared BSAlign to three striped-SIMD programs: SSW[11](version:1.0), parasail[19](version:2.4.3), ksw2[15](version:current), WFA[17](version:v2.2) and BA[16](version:0.2.0). Note that ksw2 implemented the difference recurrence relation[12] and was a component of minimap2[15]. The scores for the match, mismatch, gap open, and gap extension were set at 2, -4, -4, -2 for all implementations, respectively. For “edit mode”, we compared BSAlign to Myers[20](version:myersagrep) and Edlib[21](version:1.2.6). BA was run by rust WASM 128 bits; block size range from 32 to 2048 bp. Myers’s bit-vector algorithm was one of the fastest deterministic alignment algorithms, but it did not support global alignment and did not trace back the optimal path. Edlib extended Myer’s bit-vector algorithm with additional methods and traced back the optimal path.

We use the same real datasets as BA[16]. The short read dataset is 100,000 pairs of 101 bps Illumina HiSeq 2000 reads(accession number ERX069505). The long read dataset is 12,477 pairs of around 1000 bps Oxford Nanopore MinION reads(accession numbers ERR3278877 to ERR3278886).

We simulated query sequences and reference sequences following a configuration that approximated the error rate in the real dataset for benchmarking. We randomly selected 100 start positions that contained no gap within a 100 kb region from GRCh38. We benchmarked software in three ways: time for length, time and accuracy for divergence and length, and accuracy for long indel and band width. In the time to length comparison, we set the reference sizes as 10^2^, 10^2.25^, 10^2.5^, 10^2.75^, 10^3^, 10^3.25^, 10^3.5^, 10^3.75^, 10^4^, 10^4.25^, 10^4.5^, 10^4.75^ and 10^5^ base pairs with rounding. Then, we use PbSim2[22] to simulate a query sequence for each reference region. These query sequences and their reference sequences became pairs of input data in pairwise alignment. Sequences were simulated for both Pacbio and Nanopore using hmm model P6C4 and R103. The similarity and mutation ratio (in the format of substitution:insertion:deletion) are set as default value in PbSim2 (85% and PacBio 6:50:54, Nanopore 23:31:46). In the time and accuracy for different divergence and length comparison, we set the reference size as 10^2^, 10^3^, 10^4^ and 10^5^ base pairs. For each reference size, we also simulated reads with difference divergence(80%, 95% and 99%). In the accuracy to long indel and band width, the reference size is 10^4^ base pairs and the divergence is 80%. We randomly inserted or deleted 50, 100 and 200 base pairs sequences in the middle of the reference. For each indel size, we benchmarked software with different band width sizes (32, 64, 128, 256, 512 and 1024 base pairs).

## 4. Results

We developed the above algorithms under x86 processors using AVX2 SIMD and tested these programs on a machine with an AMD EPYC 7H12 processor, 1TB RAM, and Ubuntu Linux 20.04.1. The execution time was calculated as the sum of user and system time in a single thread. We repeated each alignment experiment 1000 times in repeat mode. To achieve a fair comparison, we modified the implementations to add a repeat mode in SSW[11] and Myers[20]. However, we were unable to add a repeat mode for parasail[19]. We developed a standard Needleman-Wunsch implementation to evaluate the alignment accuracy. SSW[11] performed local alignment instead of global alignment, we also develop a Smith-Waterman implementation to evaluate SSW’s accuracy. The recall rate was defined as the percentage of alignments that was the same score as the Needleman-Wunsch or Smith-Waterman implementation for global or local alignment, respectively. which may trim the tip sequence and get a higher score in comparison. We also recorded the maximum memory in the system during the software execution. Overall, BSAlign outperformed all the other programs or was on par with the best program in all of the experimental scenarios.

### 4.1. Evaluation on real data

Table3 showed the time and accuracy performance results for 5 algorithms evaluated using both real and simulated datasets. In the case of processing real datasets, BSAlign was the only algorithm that maintain 100% recall rate for two datasets. WFA was the fastest algorithm for Illumina reads. In “no band” mode, all algorithms aligned the whole sequences without any band width. BSAlign was 1.5-5.5X times as fast as ksw2 and SSW for Oxford Nanopore read. In “band” mode, all algorithms can align part of sequences associated with the best alignment according to its method. BA was the fastest algorithm for Oxford Nanopore reads with a recall rate of 87%. BSAlign was the second fastest algorithm with a recall rate of 100%. Overall, BSAlign was the fastest algorithm with the best recall rate for the Oxford Nanopore dataset.

**Table 3:**
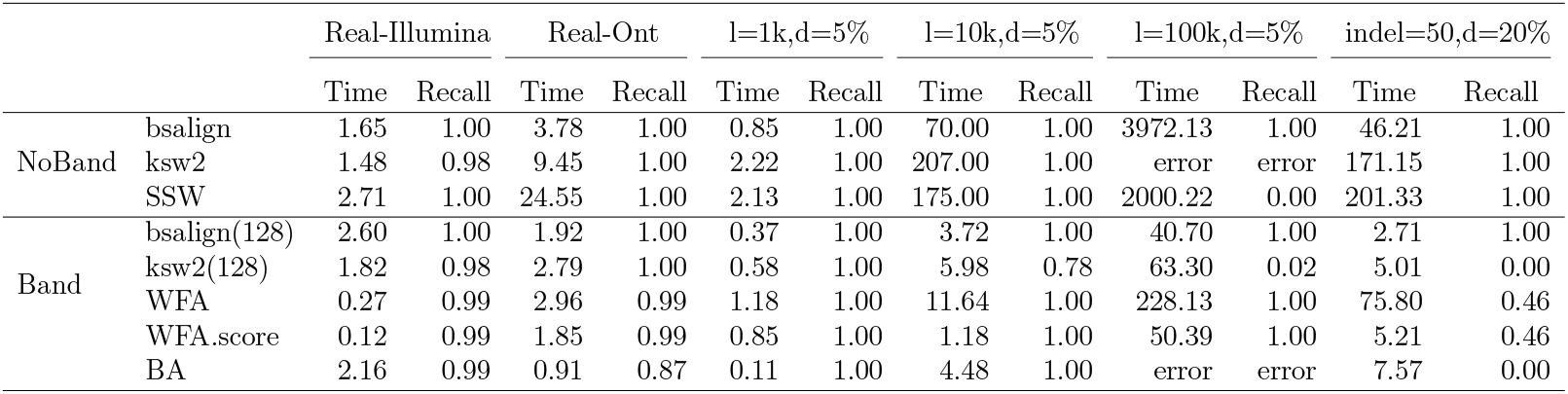
Time and accuracy performance of pairwise alignment algorithms. WFA.score only computes the alignment score, not the complete alignment.

### 4.2. Evaluation of time and accuracy for different lengths

We evaluated all software in three different ways: the running time for different lengths, the running time and accuracy for different divergences and read lengths, and finally the accuracy for different sizes of indel and different band widths. In pairwise alignment experiments, BSAlign ran faster than ksw2[15], SSW[11], and parasail[19] in both “No Band” and “Band” modes(Table 3 and Figure 3a). Among all algorithms trailed, only BSAlign and WFA[17] have the capacity to align sequences up to 100 kbps in length. Block aligner[16] was at most 3.36 times as fast as bsalign for 1,000 bps sequences. When the sequence length was equal to or longer than 10,000 bps, bsalign was at most 1.20 to 5.61 times as fast as block aligner and other algorithms.

**Figure 3:**
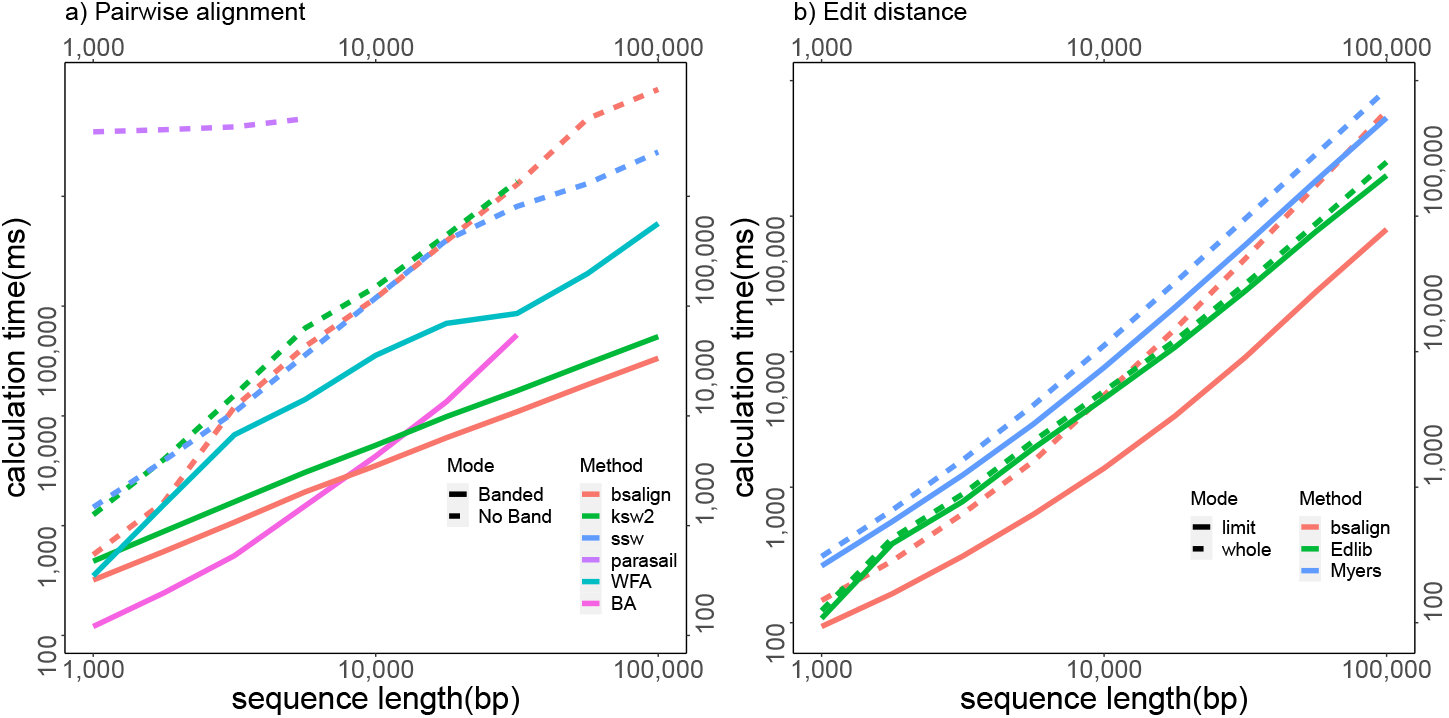
The average computation time in microseconds for Pairwise alignment and Edit distance. The length of the query sequence pair is from 1,000 to 100,000. Each pair is run 100 times. In **a**, six implementations: BSAlign, ksw2[15], SSW[11], parasail[19], wavefront alignment[17] and block aligner[16] are compared. Option band width is set at 128 for BSAlign and ksw2. In **b**, three implementations: BSAlign, Edlib[21] and Myers[20] are compared. The mode “whole” and “limit” mean the maximum edit distance is set at the whole query length and the true edit distance in simulatio, respectively.

### 4.3. Evaluation for edit distance mode

For the edit distance implementation, BSAlign recorded the fastest speed compared to Myers[20] and Edlib[21](Figure 3b). The implementations were compared in two modes. In the “whole mode”, all the aligners searched the whole sequences for the minimum edit distance. In the “limit mode”, the minimum edit distance was specified. All the aligners were instructed to stop searching the sequences that were over the minimum edit distance. In the “whole mode”, Figure3b showed that the fastest implementation switched between BSAlign and Edlib in different sequence lengths. In the “limit mode”, all the aligners ran faster than the “whole mode” due to smaller searching space, where BSAlign, Myers, and Edlib were 2.1, 1.33, and 1.11 times faster on average, respectively. BSAlign ran 2.49 and 4.93 times as fast as Myers and Edlib, respectively. Additionally, BSAlign in edit distance mode is always faster than all the pairwise alignment tools.

### 4.4. Evaluation of time and accuracy for different divergence

Furthermore, we benchmarked this six software for time and accuracy performance under different divergences (Table4). The accuracy of most software was 100%, except for the small size of band width for high divergence sequences. Most software’s time was stable in terms of processing time for different divergences except WFA[17]. Its time for high divergence sequences (20%) was 4.2 to 14.8 times slower than low divergence sequences’. When the length was 1000 bps, WFA and BA are fast and accurate. When the length increase to 10,000 bps, BSAlign(band width 128) is always fastest than other software. When the length further increased to 100,000 bps, BA(capacity overflow), parasail(early termination) and ksw2(core dumped) collapsed due to memory limitation. BSAlign(band width 128) was reliable and 6.70 times faster than other software.

**Table 4:**
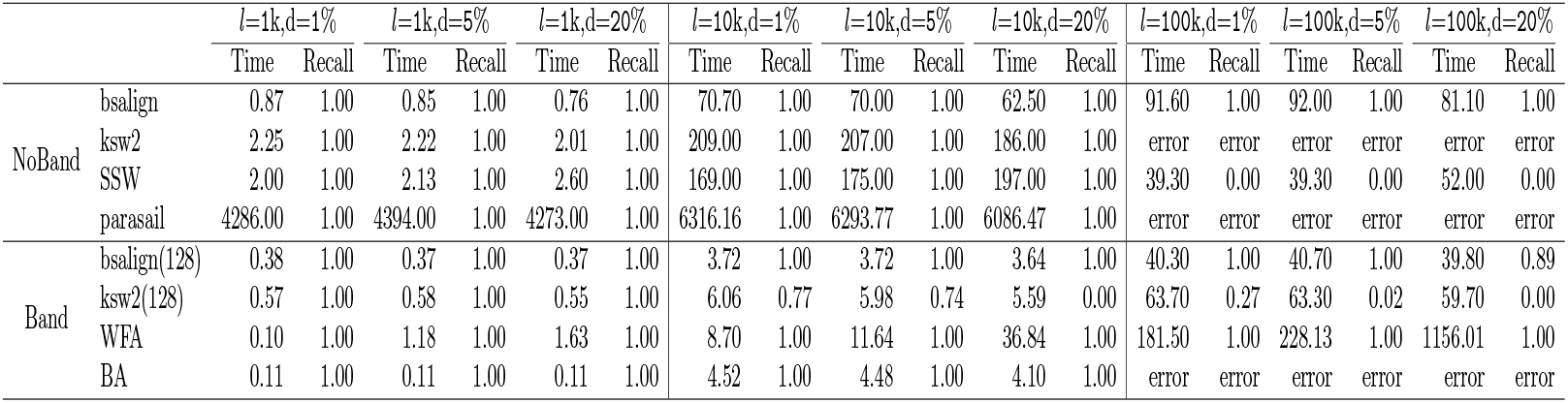
Time(ms) and accuracy performance for different divergence. *Due to the higher deletion rate in simulation, the total sequence length of 20% divergence are 4.2% and 4.6% shorter than 5% and 1% divergence’s on average, respectively.

### 4.5. Comparison of indel size, band width and accuracy

Because the banded methods might miss the optimal path, we further evaluated the influence of indel size on the alignment accuracy(Table5). In this context, we randomly inserted or deleted 50, 100 and 200 bp sequences in the middle of a reference(length=10k, divergence=20%). Overall, ksw2, SSW and BSAlign in “no band” mode were 100% correct. In band mode, WFA detected approximately 50% long indels while BA detected none. ksw2 detected all the indels when the band width was set at 1024 bps. BSAlign with band width 128, 256 and 512 bps detected all the 50, 100 and 200 indels, respectively. As expected, to accurately detect long indels, the band width size should be two times larger than the indel size. It suggests that BSAlign’s striped move is the most accurate strategy to find the optimal path. Regarding the speed, BSAlign is always faster than others. Notably, the lazy F loop implementation SSW[11] run slower for long indels while the active F loop implementation BSAlign’s runtime was stable for all the indel sizes. It suggests that BSAlign’s active F loop is fast and stable.

**Table 5:** Time(ms) and accuracy for difference indel size(50,100 and 200 bp) and band size(128 to 1024 bp).

|  |  | indel=50,d=20% |  | indel=100,d=20% |  | indel=200,d=20% |  |
| --- | --- | --- | --- | --- | --- | --- | --- |
|  |  | Time | Recall | Time | Recall | Time | Recall |
| No Band | bsalign | 46.21 | 1.00 | 45.93 | 1.00 | 45.49 | 1.00 |
|  | ksw2 | 171.15 | 1.00 | 171.04 | 1.00 | 171.00 | 1.00 |
|  | SSW | 121.90 | 0.99 | 127.88 | 0.99 | 153.19 | 0.99 |
| Band 128 bp | bsalign | 2.71 | 1.00 | 2.65 | 0.01 | 2.64 | 0.00 |
|  | ksw2 | 5.01 | 0.00 | 5.10 | 0.11 | 4.90 | 0.15 |
| Band 256 bp | bsalign | 3.26 | 1.00 | 3.30 | 1.00 | 3.22 | 0.00 |
|  | ksw2 | 9.51 | 0.43 | 9.46 | 0.42 | 9.28 | 0.38 |
| Band 512 bp | bsalign | 4.66 | 1.00 | 4.65 | 1.00 | 4.62 | 1.00 |
|  | ksw2 | 18.15 | 1.00 | 18.21 | 0.98 | 18.03 | 0.83 |
| Band 1024 bp | bsalign | 9.03 | 1.00 | 9.03 | 1.00 | 8.87 | 1.00 |
|  | ksw2 | 35.16 | 1.00 | 35.12 | 1.00 | 34.69 | 1.00 |
| Band | WFA | 75.80 | 0.46 | 75.08 | 0.51 | 76.14 | 0.61 |
|  | BA | 7.57 | 0.00 | 7.47 | 0.00 | 7.48 | 0.00 |

### 4.6. Comparison about memory

We also measured the memory during the execution(Table6). All the software except WFA required similar memory for difference divergence. The memory increased as the sequence length increased. No band method required a larger memory as they stored and calculated the whole alignment matrix. The banded method required less memory than other methods. For example, BSAlign (band width 128 bps) required 16.36 and 32.46 times less memory than WFA and BA for length 10k and divergence 20%, respectively.

**Table 6:**
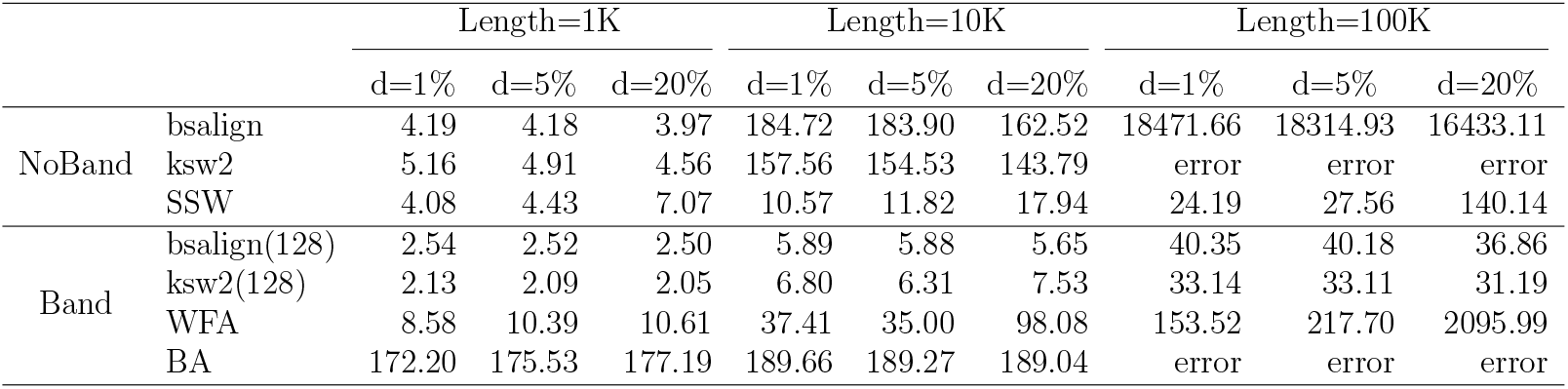
Memory (Mb) for difference size and divergence.

|  |  | Length=1K |  |  | Length=10K |  |  | Length=100K |  |  |
| --- | --- | --- | --- | --- | --- | --- | --- | --- | --- | --- |
|  |  | d=1% | d=5% | d=20% | d=1% | d=5% | d=20% | d=1% | d=5% | d=20% |
| NoBand | bsalign | 4.19 | 4.18 | 3.97 | 184.72 | 183.90 | 162.52 | 18471.66 | 18314.93 | 16433.11 |
|  | ksw2 | 5.16 | 4.91 | 4.56 | 157.56 | 154.53 | 143.79 | error | error | error |
|  | SSW | 4.08 | 4.43 | 7.07 | 10.57 | 11.82 | 17.94 | 24.19 | 27.56 | 140.14 |
| Band | bsalign(128) | 2.54 | 2.52 | 2.50 | 5.89 | 5.88 | 5.65 | 40.35 | 40.18 | 36.86 |
|  | ksw2(128) | 2.13 | 2.09 | 2.05 | 6.80 | 6.31 | 7.53 | 33.14 | 33.11 | 31.19 |
|  | WFA | 8.58 | 10.39 | 10.61 | 37.41 | 35.00 | 98.08 | 153.52 | 217.70 | 2095.99 |
|  | BA | 172.20 | 175.53 | 177.19 | 189.66 | 189.27 | 189.04 | error | error | error |

## 5. Discussion and Conclusion

Designing a dynamic banding method is about seeking a balance between speed and accuracy. No software can guarantee both fast and accurate results under any conditions. For example, WFA[17] works well for low divergence short sequences. BA[16] and ksw2[15] work well for high divergence short sequences. When the sequences are short, the band width is short as well. In this case, data structure like striped SIMD is unnecessary and time-wasting. When the sequences are long (10k bps or more), the band width increases as well. In this case, the data structure like striped SIMD is necessary and fast. The striped SIMD banded method BSAlign is faster than the anti-diagonal banding method ksw2 when the band width is 128 bps(*p* = 16, *S*_*bar*_ = 4 in 128 SSE).

## Appendix A. Availability of source code and requirements

Lists the following:

- Project name: BSAlign
- Project home page: https://github.com/ruanjue/BSAlign
- Operating system(s): Linux
- Programming language: C Language
- Other requirements: None
- License: GNU General Public License v3.0

## Appendix B. Declarations

### Appendix B.1. List of abbreviations

SW:Smith-Waterman

SIMD:single instruction multiple data

DP:dynamic programming

### Appendix B.2. List of symbols

*Q*: Query sequence.

*R*: Reference sequence.

*S*: Matching matrix.

*H*: Alignment score.

*h*: difference of H.

*E*: Alignment score end with a vertical gap.

*e*: difference of E.

*F* : Alignment score end with a horizontal gap.

*f* : difference of F.

*GapO*: gap open, less than zero.

*GapE*: gap extension, less than zero.

*GapOE*: *GapO* + *GapE*.

*S*: The number of divided segments(for whole query sequence).

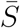: The number of divided segments(for band width).

*p*: The number of segments in a SIMD register.

*N* : register(for any value).

*M* : register in striped order(for any value).

*MF* : register in striped order(for *f*).

*i*: column id.

*j*: row id.

*u*_*i,j*_: vertical difference between H(*H*_*i,j*_ − *H*_*i−*1,*j*_).

*v*_*i,j*_: horizontal difference between H(*H*_*i,j*_ − *H*_*i,j−*1_).

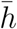: *h* in new code for edit distance mode.

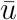: *u* in new code for edit distance mode.

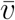: *v* in new code for edit distance mode.

### Appendix B.3. Consent for publication

Not applicable.

### Appendix B.4. Competing Interests

The authors declare that they have no competing interests.

### Appendix B.5. Funding

This study was supported by the National Key Research and Development Project Program of China (2019YFE0109600) and the National Natural Science Foundation of China (31822029 and 32200517).

### Appendix B.6. Author’s Contributions

J.R. conceived the project, designed the algorithm, and implemented BSAlign. H.S. contributed to the development and drafted the manuscript. Both authors evaluated the results and revised the manuscript.

## Appendix C. Acknowledgements

We thank Shigang Wu from CAAS for the help in the sequence alignment comparison. We thank Fan Zhang from CAAS for the help in manuscript preparation.

## Appendix A. Supplementary Figures

**Figure S1:**
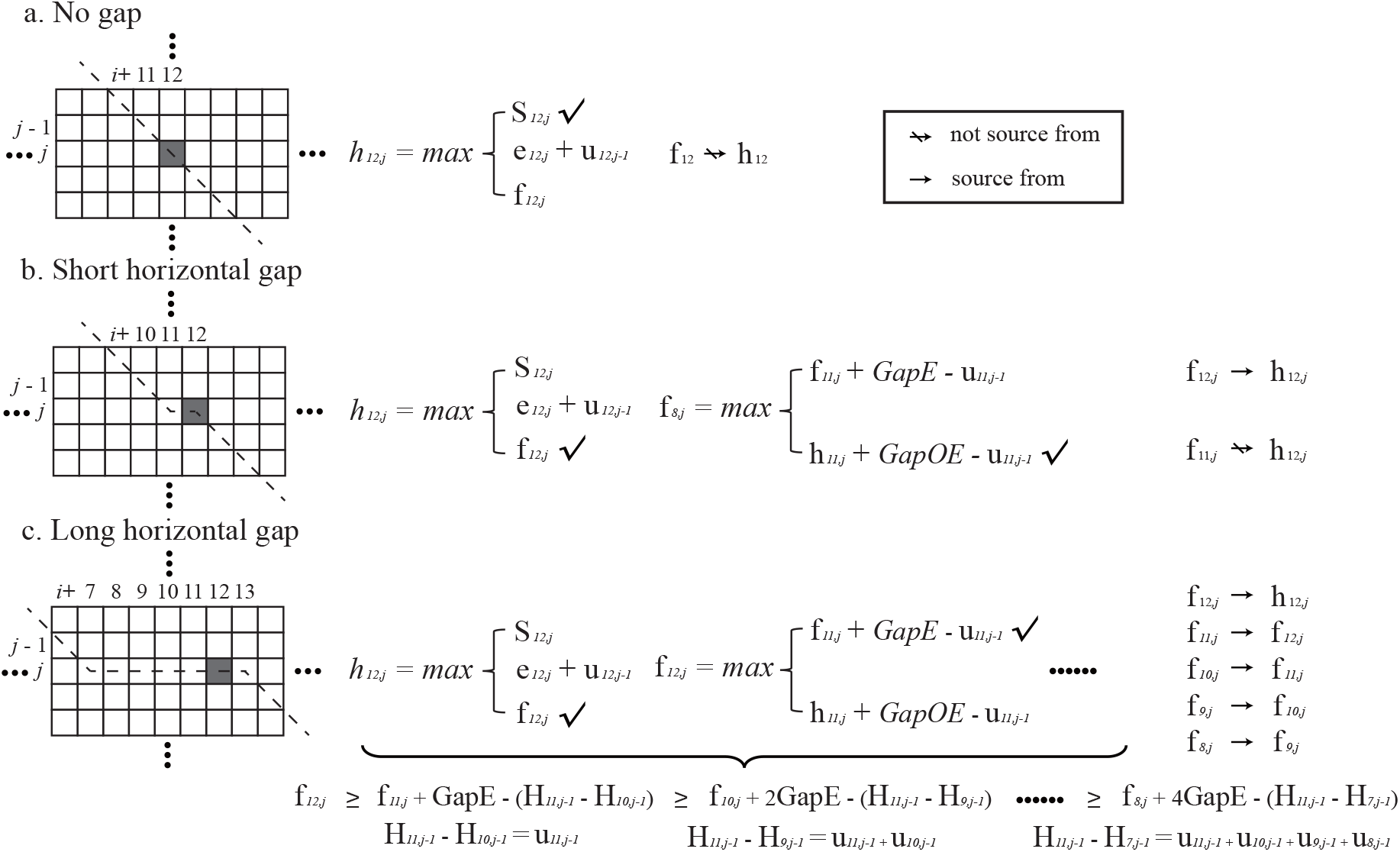
The F loop in non-striped order. (a-c) shows how the pairwise alignment calculates a cell (*h*_12,*j*_) in non-striped order in three situations: (a) no gap, (b) short horizontal gap and (c) long horizontal gap. The dash line indicates the optimal path.

